# Adaptation of motor control strategies and physiological arousal during repeated blocks of split-belt walking

**DOI:** 10.1101/2025.09.03.673831

**Authors:** Beier Lin, Kaya J. Yoshida, S. Jayne Garland, Tanya D. Ivanova, Lara A. Boyd, Courtney L. Pollock

## Abstract

Physiological arousal, mediated by the autonomic nervous system (ANS), is known to co-modulate with adaptation of motor control strategies, governed by the central nervous system (CNS), in response to repeated standing perturbations. However, adaptation of the ANS physiological arousal response during repeat exposure to walking challenges remains unknown. This study examines the physiological arousal response (electrodermal activation, EDA) and motor control strategy of gait during a single session of repeated exposure to blocks of split-belt walking. Twenty young adults completed three repeated blocks (3.5 min each) of split-belt walking (2:1 speed ratio) alternating with three blocks of tied-belt walking. Step length symmetry (SLS), EDA, bilateral tibialis anterior (TA) and gastrocnemius medialis (GM) muscle activation and ground reaction forces (GRFs) were measured. For each walking block, the first (early) and last (late) 15 strides were analyzed. A linear mixed-effects model (LMM) tested the effect of repeated blocks and phases (early/late) on SLS. Statistical parametric mapping (SPM) examined patterns of within-block changes in EDA, muscle activation, and GRFs. The greatest within-block adaptation of SLS occurred during first exposure to split-belt walking (S1; *p*<.001). Similarly, the largest magnitude of within-block adaptation in muscle activation, GRFs, and EDA occurred during S1 (*p*<.05). Attenuated EDA responses together with lower magnitude of motor adaptation in subsequent split blocks 2 and 3 were observed, indicating savings across the ANS and CNS. Taken together, these findings demonstrate that the ANS-mediated physiological arousal response modulates alongside the CNS-driven locomotor adaptation to repeated exposure to blocks of split-belt walking.

## Introduction

Human locomotion is inherently adaptable, allowing individuals to modify gait patterns to maintain stability and energy efficiency, the two primary goals of walking (Kuo & Donelan, 2010; Selinger et al., 2015). When confronted with walking perturbations, the central nervous system (CNS) rapidly identifies the error and recalibrates sensorimotor control to restore stability to the altered walking conditions; a process known as motor adaptation (Bastian, 2008; Izawa et al., 2008). A common response to instability involves increased neuromuscular activation around key joints to enhance postural control and movement correction (Krause et al., 2018; Riemann & Lephart, 2002; Torres-Oviedo & Ting, 2007). Beyond CNS-driven neuromuscular responses, challenges to balance also engage the autonomic nervous system (ANS), with the physiological arousal response shown to modulate alongside motor responses to repeated standing balance perturbations (Carpenter et al., 2006; Sibley et al., 2008). However, whether the CNS and ANS co-modulate during motor adaptation to challenging walking conditions remains unclear, leaving a critical gap in understanding motor adaptation under dynamic balance challenges.

Use of the split-belt treadmill facilitates the study of adaptation of sensorimotor control during continuous walking (Malone & Bastian, 2010; Morton & Bastian, 2006; Reisman et al., 2005). By imposing different belt speeds on each leg, split-belt walking paradigms introduce a controlled perturbation that alters gait patterns (Reisman et al., 2005). Studies have shown that individuals exhibit asymmetrical walking when first exposed to split-belt walking, but rapidly adapt to the new walking demands, achieving a more symmetrical gait pattern by the end of a single block (Bruijn et al., 2012; Malone & Bastian, 2010; Sombric & Torres-Oviedo, 2020). Electromyography (EMG) analyses during split-belt walking adaptation have shown reorganization of muscle activation patterns of the ankle, such as heightened muscle activation in tibialis anterior (TA) and gastrocnemius medialis (GM) (Dietz et al., 1994; Finley et al., 2013; MacLellan et al., 2014; Ogawa et al., 2014). MacLellan et al. (2014) further reported that muscle activation timing shifts during split-belt walking adaptation, with some activation patterns maintaining their structure while others adjusting to earlier onset in the gait cycle. Notably, these adjustments occur bilaterally, suggesting that the neuromuscular system does not regulate each limb independently but instead coordinates interlimb adaptation to maintain gait stability under altered walking conditions (MacLellan et al., 2014).

The ANS-mediated physiological arousal response has been shown to co-modulate with motor control strategies during postural control responses to standing perturbations (Carpenter et al., 2006; Critchley, 2002; Pollock et al., 2017; Sibley et al., 2008). Height-induced postural threats, such as when participants stand on an elevated platform, have been found to elicit consistent changes in motor control during quiet standing balance and heightened physiological arousal states (Horslen & Carpenter, 2011; Johnson et al., 2019). Moreover, perturbation-evoked physiological arousal, as measured by electrodermal activity (EDA), increases in conjunction with both the perturbation size and task complexity, suggesting that physiological arousal levels scale with postural challenges (Sibley et al., 2010a). Yet, little is known about the ANS-mediated physiological arousal in modulating motor control during continuous locomotor adaptation, such as walking.

This study aims to investigate adaptation of motor control strategies and the physiological arousal response during repeated exposures to perturbations induced by split-belt walking. Specifically, we explore how CNS-driven motor adaptation co-modulates with the ANS-mediated physiological arousal response over three blocks of split-belt walking (i.e., adaptation) interspersed with washout blocks of tied-belt walking (i.e., de-adaptation) in a single session. We hypothesize that: 1) the initial block of split-belt walking will demonstrate the greatest magnitude of locomotor adaptation and attenuation of the physiological arousal response by the late phase of the block; and 2) repeated exposures to split-belt walking perturbations will demonstrate smaller magnitudes of within-block motor adaptation and physiological arousal response, suggestive of motor savings and an attenuation of the physiological arousal response.

## Methods

### Subjects

Twenty young participants (aged 18-35 years old; 10 females, 10 males) naïve to split-belt treadmill walking completed the study. Inclusion criteria were no history of neurological or musculoskeletal impairments affecting gait or walking function and no history of depression or anxiety requiring medication. All participants provided written consent before participation. The study adhered to the latest revision of the *Declaration of Helsinki* and received approval from the University of British Columbia Clinical Research Ethics Board (H21-02171).

### Experimental protocol

All participants completed the 10-Meter Walk Test (10MWT), performing two trials each at comfortable gait speed and fast gait speed (FGS). The average FGS across two trials was used to set individualized treadmill speeds for the split-belt walking protocol. The fast belt speed was set to 90% of participant’s average FGS while the slow belt speed was 50% of the fast belt speed to maintain a 2:1 ratio between the belt speeds. The fast belt was always assigned to the left side for all participants.

Participants then completed a 32.5-minute treadmill walking protocol (M-GAIT, MOTEK Medical) with an overhead safety harness (Fig 1a). The harness did not support body weight or interfere with normal walking gait patterns. Handrails were also available on both sides of the treadmill; participants were encouraged not to use them unless necessary. Participants were familiarized with the treadmill speeds, and the baseline walking patterns were established. This familiarization was followed by alternating 3.5-minute blocks of split-belt (belts moving with 2:1 ratio between the belt speeds) and tied-belt (equal slow belt speeds) walking (Fig 1b). Participants were instructed not to talk during data collection and avoid looking down at the belts to gain any visual feedback about the belt conditions.

**Figure 1.**
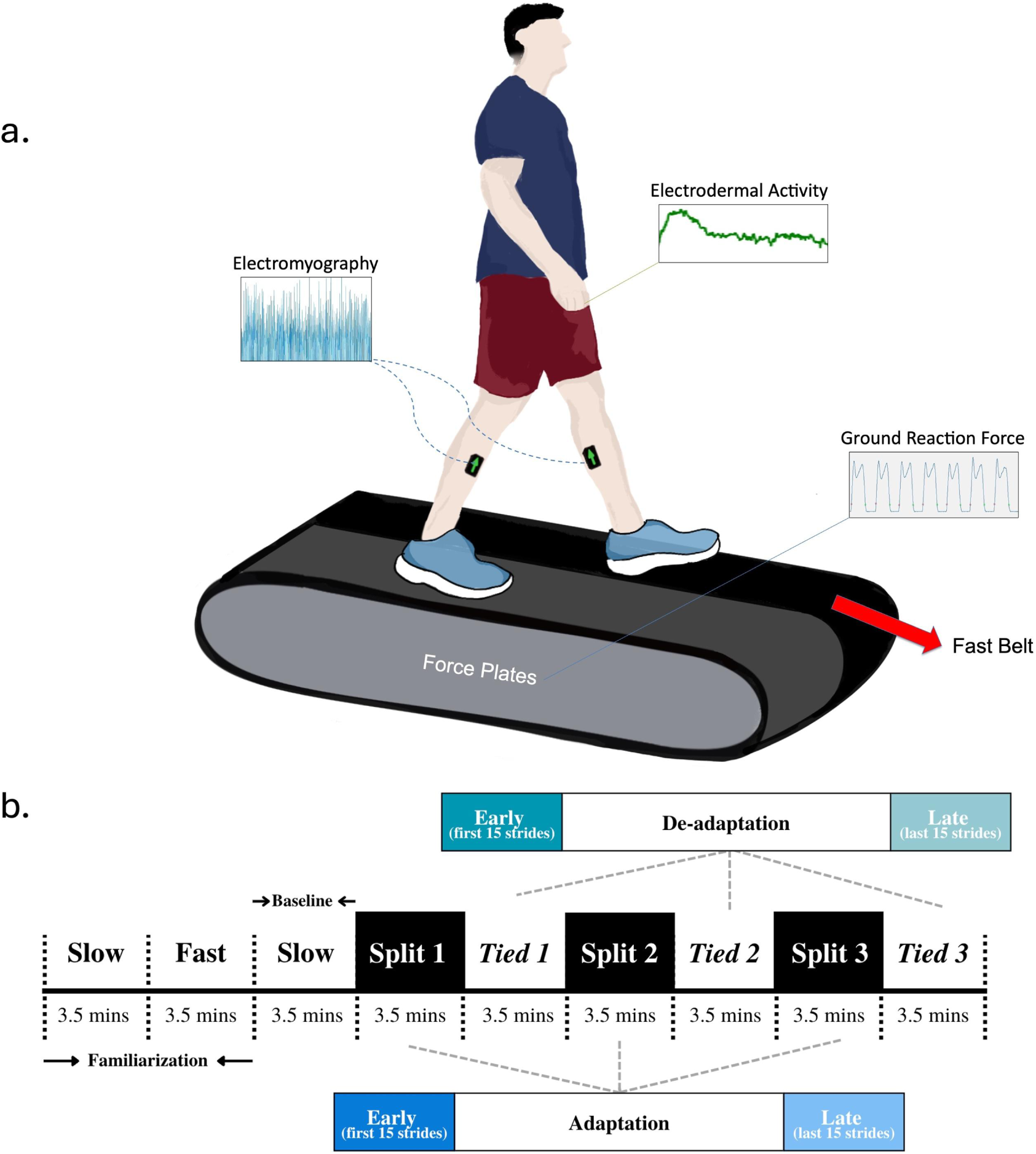
Experimental protocol during treadmill walking. A) Electromyography (EMG) was measured bilaterally from TA and GM; electrodermal activation (EDA) was measured from the palmar surface of the hand; ground reaction forces (GRF) were recorded by the treadmill embedded force plates under each belt. Participants were not informed about the change of belt conditions. B) Timeline of the experimental protocol. The protocol started with 2 familiarization blocks of tied-belt walking at slow and fast speed, respectively, followed by a block of slow speed used for baseline measure. Then the split-belt adaptation blocks (shown with black background) alternated with the tied-belt de-adaptation blocks followed. The duration of each block was 210s (3.5 mins). The Early and Late phase in each block of walking was defined as the first 15 strides (excluding the first step) and last 15 strides within each block.

### Measures

#### Electromyography (EMG)

EMG recordings of bilateral tibialis anterior and gastrocnemius medialis were collected using Trigno wireless surface electromyography (EMG) sampled at 2,184 Hz and EMGworks to capture adaptation of muscle activation of the ankle (Delsys, Inc., Natick, MA, USA). Prior to electrode placement, the skin surface was cleansed with alcohol pads. EMG signals were pre-processed in MATLAB (Mathworks_R2023a, Natick, MA) as follows: detrended, band-pass filtered (20-450 Hz, 4^th^-order Butterworth), full-wave rectified, and root mean square was calculated over a 50 ms window to obtain smoothed envelopes. Processed EMG signals were normalized to the mean activation over a continuous 15 stance phase during slow speed tied-belt baseline walking (Fig1b, baseline).

#### Electrodermal activity (EDA)

Physiological arousal was collected using electrodermal activity (EDA, Cambridge Electronics Design LTD, Cambridge, UK) from the palmar surface of the right hand at 300 Hz. Bipolar electrodes were positioned on the hypothenar and thenar eminences (Boucsein, 2012). EDA signals were zero-phase low-pass filtered at 10Hz, followed by median filter with a 500ms window to eliminate motion artifacts (Barua et al., 2020). After pre-processing, EDA data were baseline corrected to the average of a plateau period from baseline walking. EDA data were then normalized to the within-session peak value (presented as percentage max, %max).

#### Ground reaction forces (GRFs) and step length symmetry (SLS)

Ground reaction forces (GRFs) were recorded from embedded treadmill force plates to measure the changes in kinetic forces generated during adaptation to split-belt walking. All GRFs were low-pass filtered at 10Hz. Step length was determined by the anterior-posterior distance between the center of pressure for each foot at the onset of double stance phase, defined as the moment when a vertical GRF greater than 15% body weight on both sides.

Step length symmetry (SLS), an indicator of locomotor adaptation during split-belt walking, was calculated as a normalized difference between step lengths on the fast and slow belts, as outlined in Equation 1 (Malone & Bastian, 2010). An SLS value of zero indicates perfect step symmetry. Positive values suggest a longer step on the fast belt, while negative values indicate a longer step on the slow belt (Reisman et al., 2010). Normalizing step symmetry to each participant’s step length allowed comparisons across individuals with different baseline step lengths.

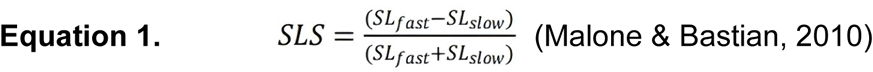

#### Rate of Perceived Stability (RPS)

The rate of perceived stability (RPS) is a numerical scale with descriptive anchors that provided a validated measure of self-perceived balance challenge during walking. At the end of the study, participants self-reported their RPS scores at the onset of each split block (Fig 2) (Espy et al., 2017).

**Figure 2.**
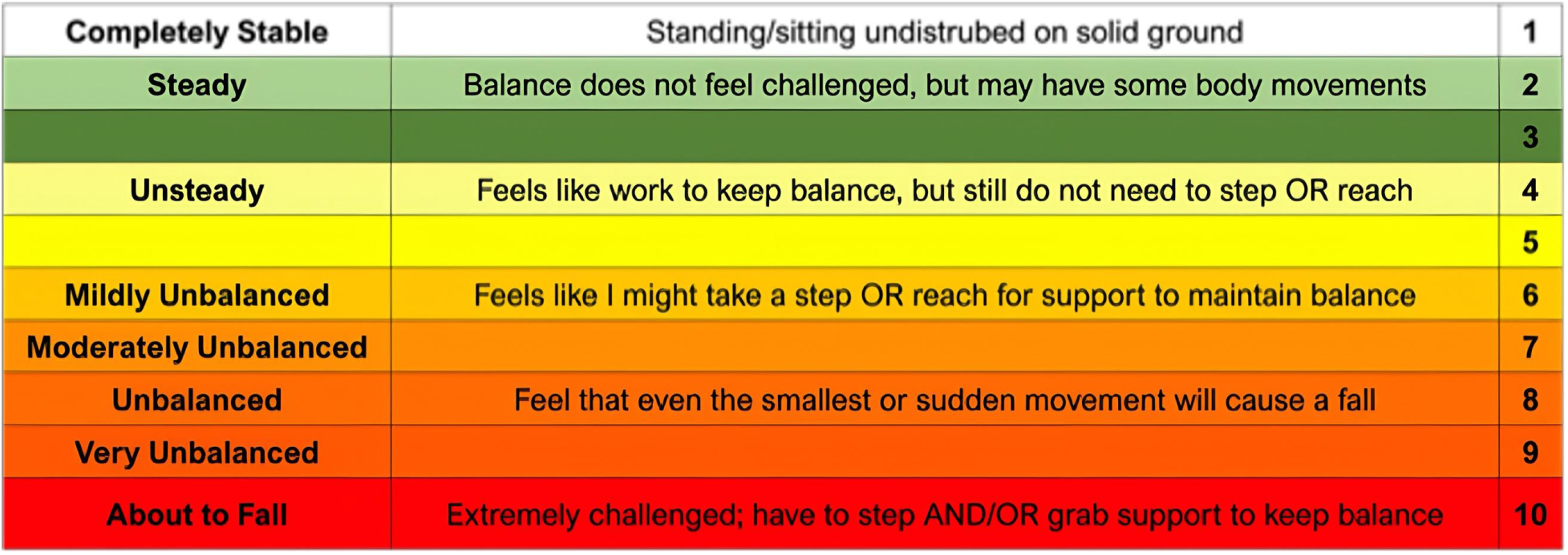
Rate of Perceived Stability (RPS) scale. All participants verbally reported their RPS score regarding each onset of the split-belt walking at the end of the study. Note: Adapted from “Intensity of Balance Task Intensity, as Measured by the Rate of Perceived Stability, is Independent of Physical Exertion as Measured by Heart Rate”, by Espy, D., Reinthal, A., & Meisel, S., 2017, Journal of Novel Physiotherapies, 7(72), Article 72. Copyright 2017 by Journal of Novel Physiotherapies.

#### Data analysis

Signals were synchronized during data collection by pulses generated by the split-belt treadmill at the initiation of each block. All signals were time normalized (EDA = 0-100% time cycle; EMG and GRFs = 0-100% stance cycle) for within block temporal analysis. For all measures, data were segmented into “Early” and “Late” phases for each of the three Split (adaptation) blocks—S1, S2, S3; and three Tied (de-adaptation) blocks—T1, T2, T3. The Early phase was defined as the first 15 recorded strides, excluding the first step to eliminate any anomalies due to the acceleration or deceleration (Alexander et al., 2005; Malone & Bastian, 2010; Ogawa et al., 2014). The Late phase encompassed the final 15 strides of each block. Baseline walking was established as the late phase (last 15 strides) of both fast and slow speed baseline tied-belt walking blocks. EMG, EDA, and GRFs were analyzed and presented as baseline data specific to belt speeds.

EDA data from one participant were removed due to poor signal quality from electrode detachment. Bilateral GM data from one participant were excluded due to sensor detachment. Additionally, one participant’s baseline walking EMG data were removed due to poor signal quality.

#### Statistical Analysis

Descriptive statistics were performed for participants’ demographic data and treadmill walking speeds. Linear mixed-effects models (LMMs) were performed to examine the effects of repeated Split and Tied blocks on SLS from the Early and Late phases of each block (R Studio 4.3.1, Package *lme4* version 1.1-34). Participant ID was included as a random effect to account for repeated measures. Fixed effects included Block (Split/Tied), Phase (Early/Late), and their interaction (model: *SLS ∼ Phase + Block + Phase*Block* + (1|Participant)). Post-hoc comparisons were adjusted using Bonferroni’s correction. Dot plots were generated from model estimates using *ggplot2* package (version 4.2-2, RStudio). A separate LMM assessed changes in RPS scores across Split blocks (model: *RPS Score ∼ Block* + (1|Participant)).

EMG, EDA, and GRF data were analyzed with statistical parametric mapping (SPM) paired t-tests for each Block (Split/Tied), with Phase (Early vs. Late) as a within-subjects factor. SPM analyses were performed in Python with open-source package *spm1d* (version 0.4.18, https://spm1d.org/). These calculations were based on a threshold value estimated by SPM {t} distribution, with the null hypothesis being rejected when the observed t-value trajectory exceeded the critical threshold corresponding to an alpha level of 0.05 (Pataky, 2010, 2012; Pataky et al., 2016). Significances were accepted at *p* < .05 for all statistical tests.

## Results

Twenty young adults (26.8 ± 3.3 years, 10 F, 10 M) participated in this study. Table 1 summarizes participant demographics and treadmill walking speeds. Self-reported perception of stability (Fig 3) was significantly higher in S1 than S2 and S3 (*p*<.001), and S2 was significantly higher than S3 (*p*=.0026).

**Figure 3.**
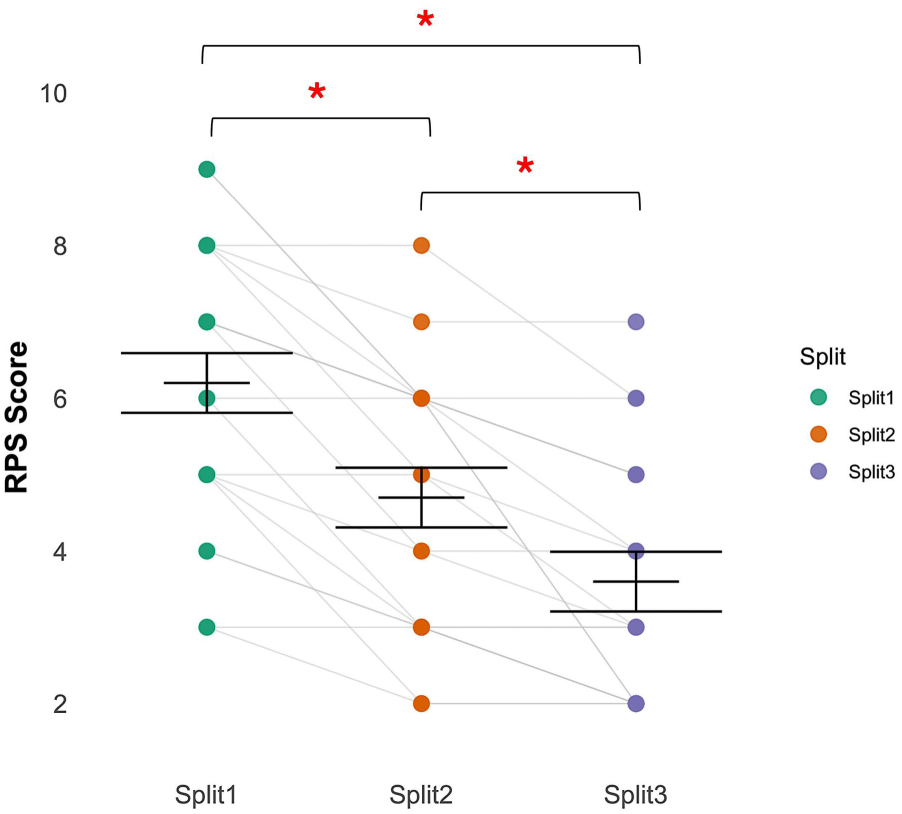
Dot plots showing differences in Rate of Perceived Stability scores across three split blocks for all participants. Error bars represent the fit for the model ± 95% CI. **p <.001; *p<.05.

**Table 1.**
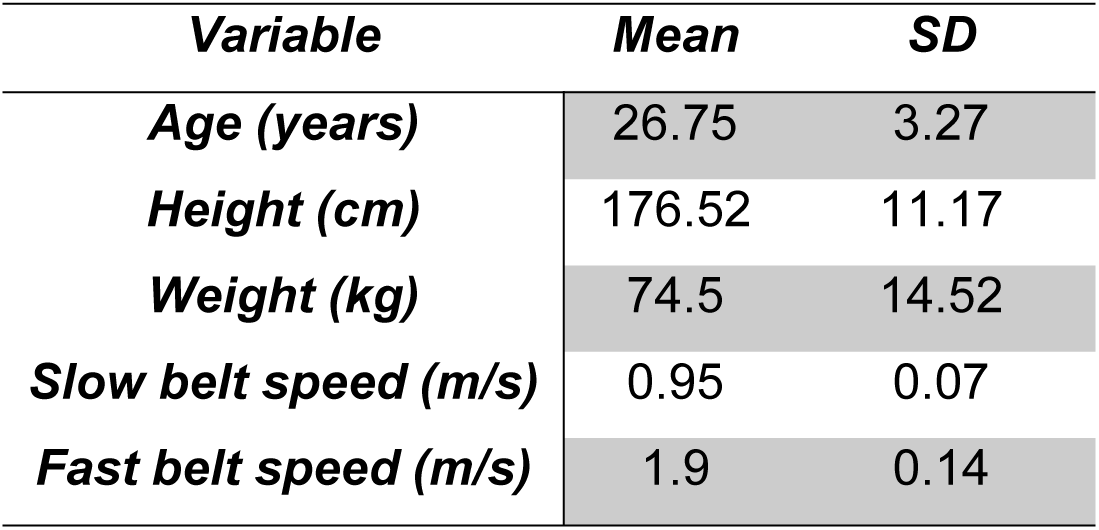
Summary of participant demographic data, walking speeds and participant reported Rate of Perceived (RPS) scores for each split block during three blocks of split-belt walking, mean and standard deviation (SD).

### Step Length Symmetry

Baseline SLS during the baseline slow tied-belt walking block was indicative of symmetrical walking among all participants (0.023 ± 0.033). Fig 4 illustrates a representative participant’s continuous step length, SLS, and EDA responses throughout the entire walking protocol. The largest difference in SLS ratio was observed during the Early phase of Split 1, coinciding with a peak in EDA signals.

**Figure 4.**
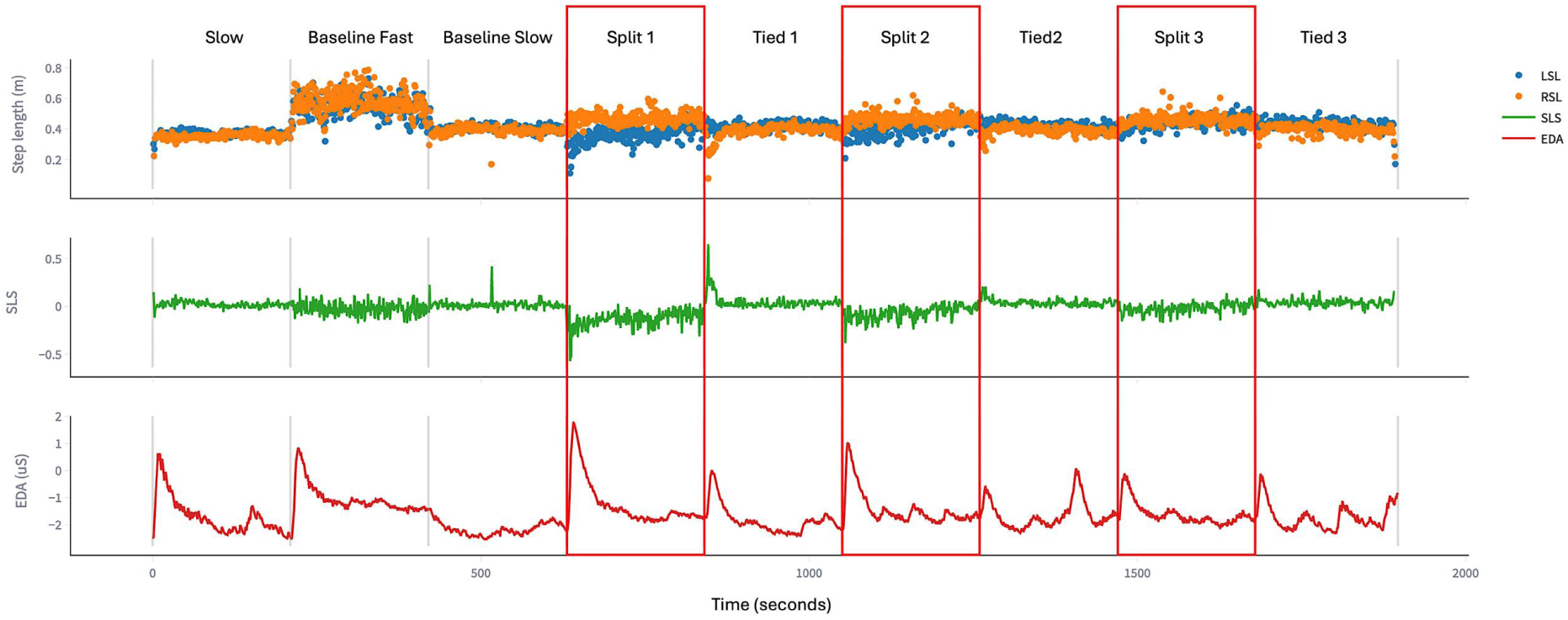
Representative raw data from a participant across the full walking trial. Top row: raw step length for the left (blue) and right (orange) foot. Middle row: Step length symmetry (SLS) ratio calculated from raw data. Bottom row: raw electrodermal activation (EDA) signal. Red boxes highlight split-belt walking blocks; unshaded regions represent tied-belt walking. During the baseline slow speed tied-belt walking before Split 1, SLS indicates symmetrical gait. The greatest step length asymmetry occurred during the Early phase (first 15 strides) of S1, coinciding with the peak EDA response across the session. SLS increased in the Early phases of S2 and S3 with EDA response attenuated alongside.

Split 1 demonstrated a significant increase in SLS from the Early to the Late phase (mean: Early = −0.27 to Late = −0.08; *p*<.001, Fig 5). Split 2 also showed significant locomotor adaptation from Early to Late phase (mean: −0.11 to −0.06; *p*=0.048, Fig 5). Comparisons of SLS during the Early phase across split blocks showed SLS in Early Split 2 and Early Split 3 were significantly more symmetrical than SLS in Early Split 1 (*p*<.001), suggestive of motor savings between blocks. During de-adaptation (tied-belt walking), the reduction in step length asymmetry showed a significant difference between the Early and Late phases of Tied 1 and Tied 2 (Fig. 5, *p*<.001). By the Late phase of each tied block, the mean SLS reached 0.03, within the range of the baseline symmetrical walking (0.023 ± 0.033).

**Figure 5.**
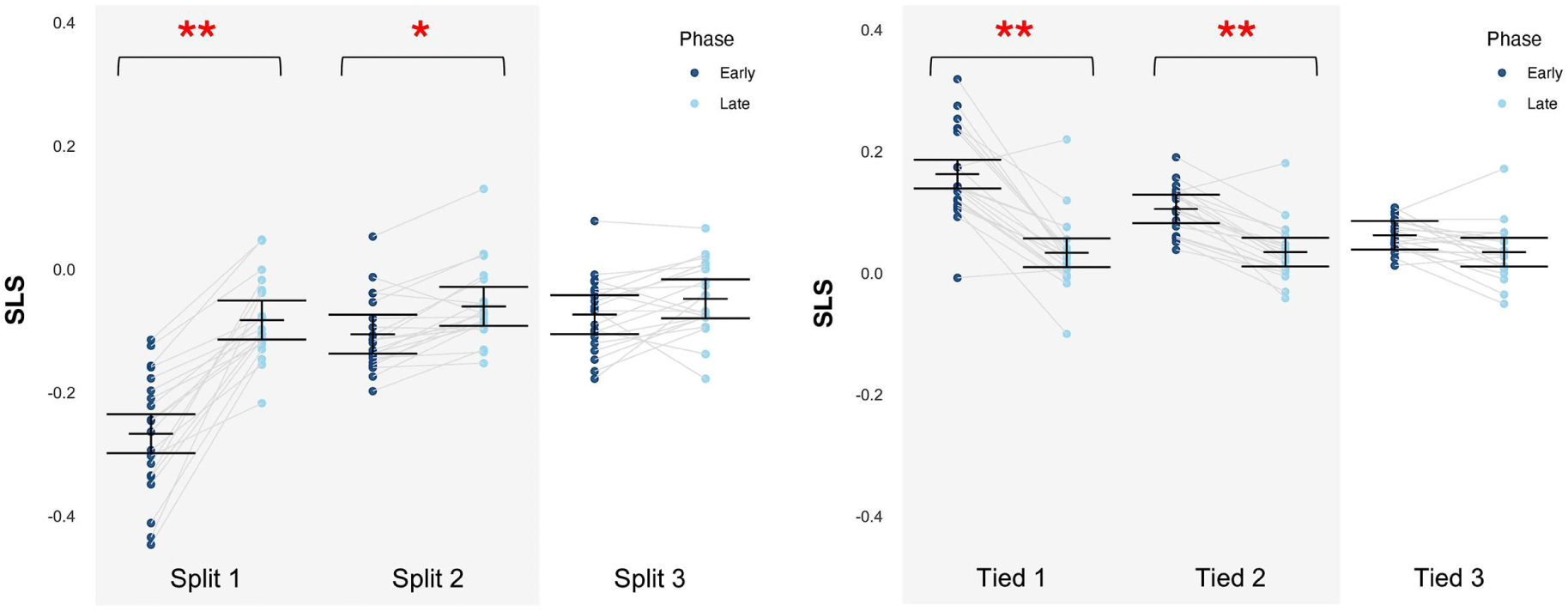
Dot plots showing differences in SLS for Early and Late phases between Splits (left) and Tied (right) across three blocks for all participants. Error bars represent the fit for the model ± 95% CI. Grey block represents statistically significant changes between early and late phases of split blocks 1 and 2, tied blocks 1 and 2. **p<.001; *p <.05.

### SPM analysis of Physiological Arousal (EDA)

SPM analysis revealed significant within-block adaptation of physiological arousal across all Split and Tied blocks, with higher EDA responses during the Early phases compared to Late phases (*p*<.05 for all; Fig 6). During adaptation split-blocks, Early phase EDA responses were significantly elevated in all Split blocks, with the highest level of physiological arousal observed during the initial exposure to split-belt walking (S1). Significant adaptation was identified comparing the Early to Late phase of S1 (*p*<.05). While Early to Late phase differences remained significant in subsequent split blocks (S2 and S3; *p<*.05), the magnitude of these differences progressively declined, suggesting habituation of arousal responses with repeated exposure to split-belt walking. Notably, late phase physiological arousal levels in each Split block converged toward that observed during fast speed tied-belt walking, approximating 20% of the maximum EDA response.

**Figure 6.**
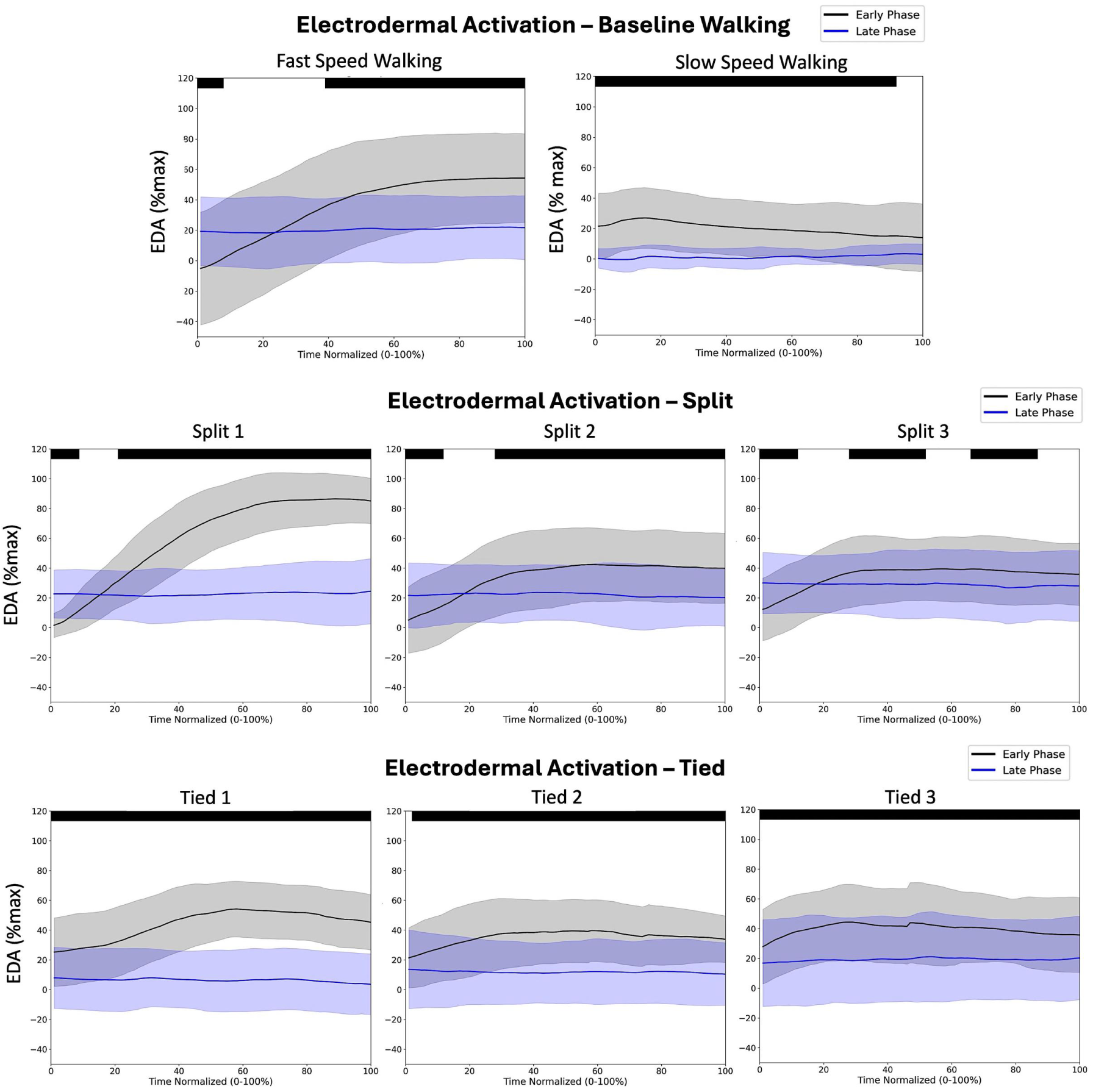
Patterns of physiological arousal presented by electrodermal activation (EDA) data during walking in 100%-time cycle respectively for the Early (black) and Late (blue) phases. Black bars on top indicate regions with statistically significant differences between phases with magnitude above the relevance criterion of t-values of the SPM. Baseline walking differences are reflective of speed change from slow to fast speed walking and from fast to slow speed walking, respectively.

During de-adaptation tied-belt blocks, EDA responses were elevated in the Early phases upon returning to tied-belt walking but steadily declined across the normalized time cycle in all three Tied blocks (*p*<.05), approaching the levels of physiological arousal observed during baseline slow speed walking.

### SPM analysis of Electromyography

Figure 7 presents the results of SPM paired t-tests on the temporal projections of the phases (Early vs. Late) during stance across blocks. Baseline walking patterns for each leg are presented specific to either the fast or slow speed assigned to that leg during split-belt walking. During split-block adaptation blocks, S1 resulted in increased GM activity in the fast leg through weight acceptance to mid-stance, while TA of the fast leg demonstrated a pattern of increase amplitude throughout stance. In the slow leg, S1 resulted in increased activation of TA throughout the stance phase with a notable early peak in activity during weight acceptance to mid-stance, while GM activity increased during push-off in late stance. Significant within-block adaptation (Early vs. Late) of bilateral activity of GM and TA (fast and slow leg) muscles during stance was observed in each of the three Split blocks (*p*< .05, respectively). However, the magnitude of these adaptations and between-phase significances in percentage of stance decreased across blocks, suggestive of motor savings with repeated exposure to split-belt walking.

**Figure 7.**
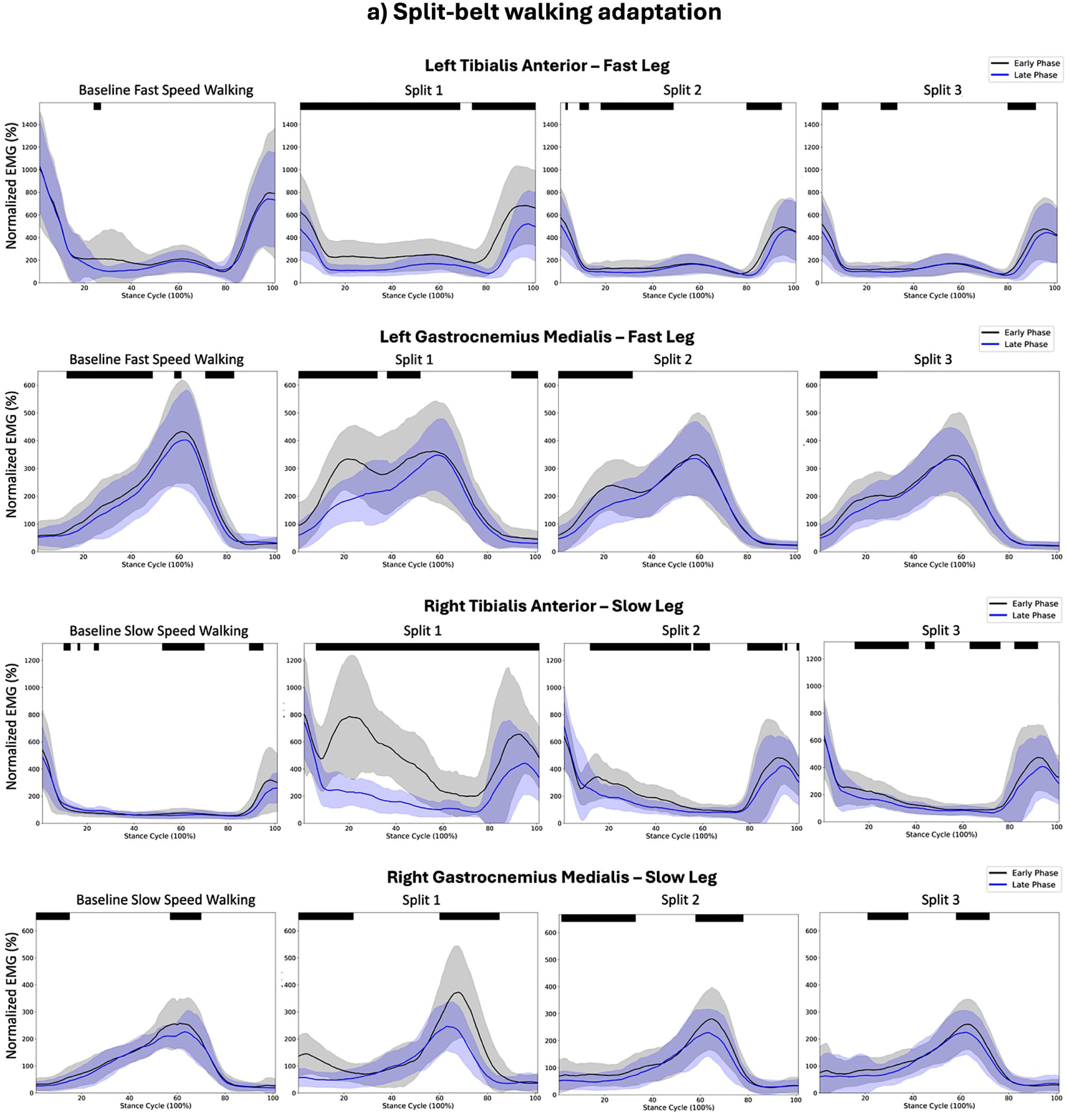
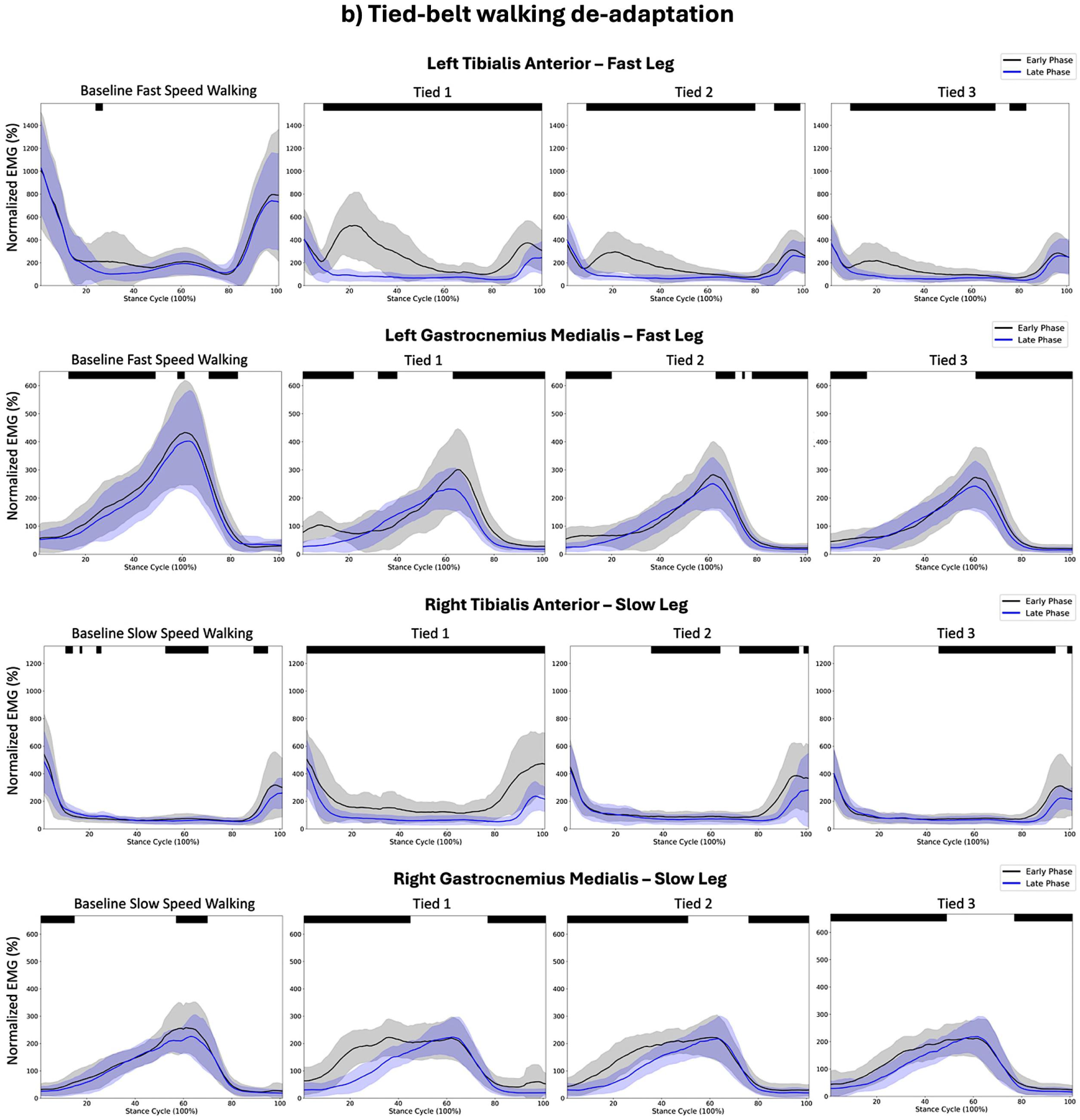
Mean (1 standard deviation) EMG patterns in 100%-time cycle respectively for the Early (black) and Late (blue) phases during stance. Black bars on top indicate regions with statistically significant differences between Early and Late phases above the relevance criterion of t-values of the SPM. A) Split-belt walking adaptation; b) Tied-belt walking de-adaptation. Baseline EMG walking data are included for each leg at the speed corresponding to its assigned belt speed during split-belt walking (i.e., baseline fast walking is shown for the fast leg; baseline slow walking is shown for the slow leg).

During tied-belt de-adaptation blocks, the fast leg exhibited elevated TA activity throughout the stance phase with an earlier onset of activation during initial stance upon return to tied-belt walking. GM activity in the fast leg also increased during late stance. In the slow leg, early-phase TA activity remained elevated across the entire stance phase in all three Tied blocks, and GM activity increased significantly from initial to mid stance. Significant within-block adaptation patterns were evident in bilateral TA and GM muscles across all Tied blocks (*p*< .05, respectively). However, similar to the Split blocks, the magnitude of these within-block changes attenuated in T2 and T3 relative to T1.

### SPM analysis of Ground Reaction Forces

SPM analysis revealed significant bilateral adaptation patterns in both the vertical (vGRF) and anterior-posterior GRF (APGRF) between the Early and Late phases of each Split and Tied block (Fig 8). During S1, the fast leg exhibited lower peak vGRF and APGRF during both initial stance and terminal stance in the Early phase compared to the Late phase (*p*<.05). In contrast, the slow leg showed a higher peak vGRF and APGRF during initial stance in the early phase (*p*<.05), followed by a gradual decline through mid- and terminal stance (*p*<.05, respectively), compared to the Late phase. This pattern persisted in S2 and S3, though with reduced magnitude of change, suggesting motor savings with repeated exposures to split-belt walking. During the de-adaptation phase, the GRF patterns reversed relative to the adaptation phase, reflecting a realignment toward symmetrical gait mechanics.

**Figure 8.**
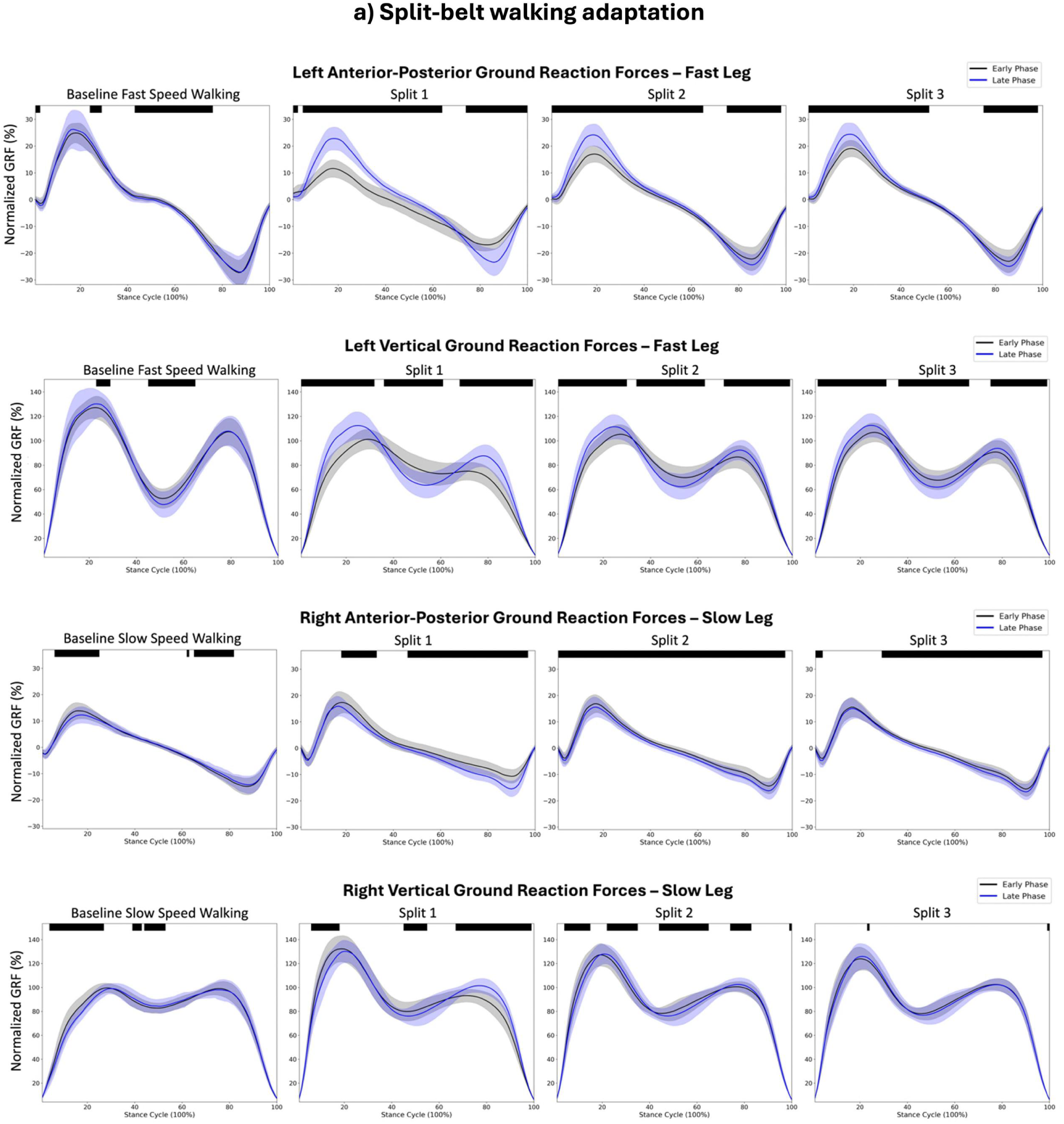
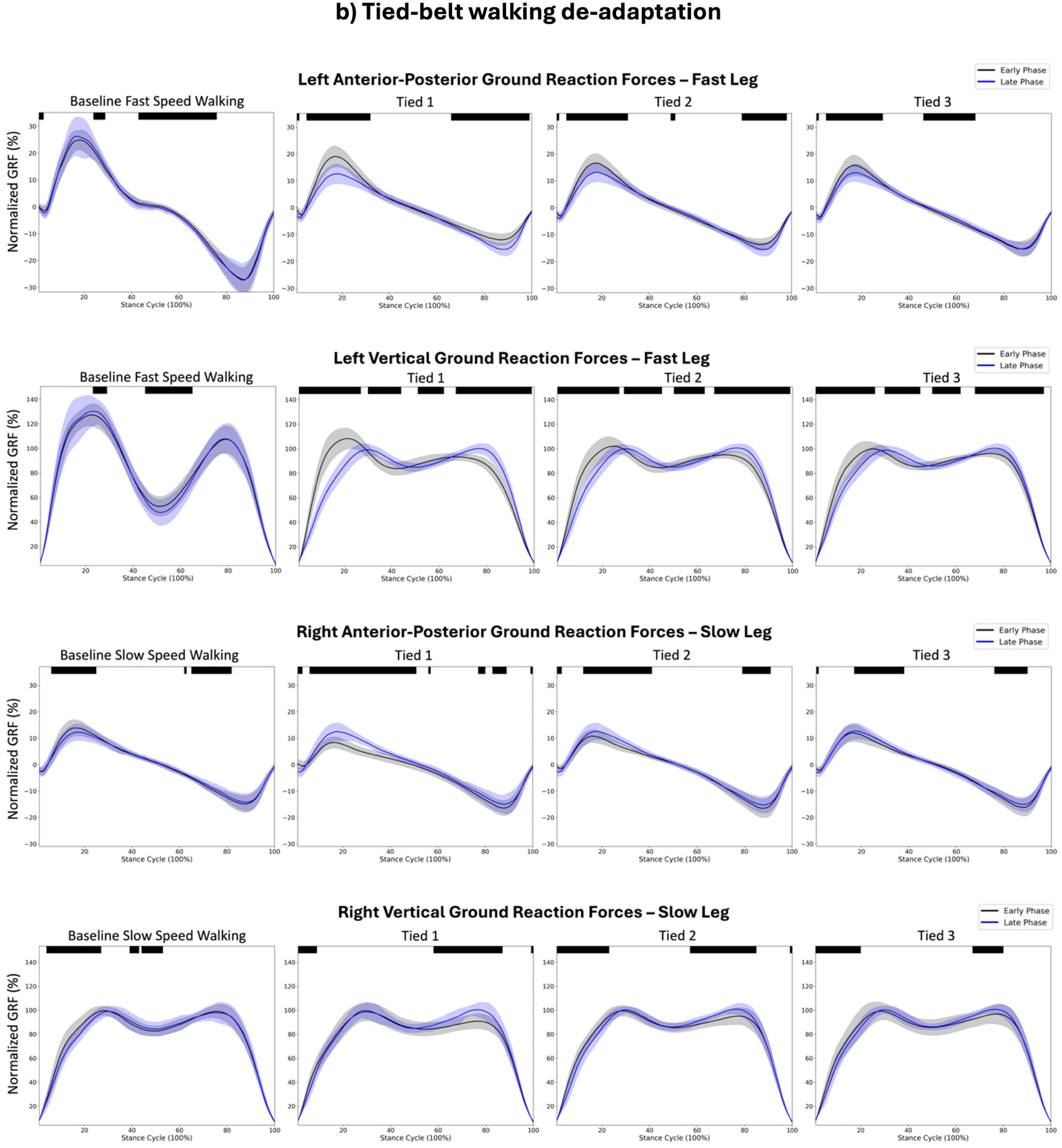
Mean (1 standard deviation) GRF patterns in 100%-time cycle respectively for the Early (black) and Late (blue) phases during stance. Black bars on top indicate regions with statistically significant differences between Early and Late phases above the relevance criterion of t-values of the SPM. A) Split-belt walking adaptation; b) Tied-belt walking de-adaptation. Baseline GRF walking data are included for each leg at the speed corresponding to its assigned belt speed during split-belt walking (i.e., baseline fast walking is shown for the fast leg; baseline slow walking is shown for the slow leg).

## Discussion

The present study explored the co-modulation of motor responses (CNS) and the physiological arousal response (ANS) during repeated exposure to blocks of split-belt treadmill walking. We found a consistent pattern of motor adaptation to split-belt walking, measured by SLS, muscle activations around the ankle, and vertical and anterior-posterior GRFs, together with physiological arousal responses across three separate practice blocks. The most pronounced within-block adaptation of SLS occurred during the first exposure (S1) to split-belt walking with clear evidence of motor savings of SLS in subsequent exposures (S2 and S3). Similarly, the magnitude of within-block adaptation pattern changes in muscle activation, GRFs, and physiological arousal was greatest during S1; savings were also observed with attenuated responses in later blocks across these measures. Overall, our results provide evidence that the CNS-driven locomotor adaptation co-modulates with the ANS-mediated physiological arousal during repeated exposure to split-belt walking, with the initial exposure eliciting the greatest magnitude of within-block adaptation, followed by motor savings evident across measures in subsequent blocks of exposure to split-belt walking.

Adaptation to split-belt walking reflects the CNS’s capacity to recalibrate locomotor control in response to altered dynamic walking demands imposed by the asymmetrical belt speeds, most often quantified by changes in between-leg SLS. Here, we showed that the SLS adaptation was most evident in the first exposure to the split-belt walking. The pattern of adaptation during S1 aligns with prior findings of motor adaptation in young adults during a single bout of split-belt walking (Malone et al., 2011; Malone & Bastian, 2010; Morton & Bastian, 2006; Reisman et al., 2005). Notably in our study, participants rated S1 as the most challenging of the three Split blocks, suggesting that the novelty and destabilizing nature of the task was most prominent in this first exposure. Aligned with participant ratings, the greatest magnitude of adaptation of walking pattern and of physiological arousal response both occurred during the first exposure. In fact, physiological arousal response decreased by approximately 80% from early to late phases, returning to a level similar to that of baseline fast speed walking (approximately 20% of session maximum) by the late phase of S1. The elevated initial arousal response is consistent with previous work exploring physiological arousal response under conditions of repeated exposure to postural threat (Horslen & Carpenter, 2011; Sibley et al., 2010b); our findings extend this understanding into dynamic locomotor adaptation.

Use of SPM analyses provides further insight into how motor and physiological arousal responses adapt during exposure to split-belt walking. Unlike discrete point analyses, SPM analyzes the entire time series and identifies statistically significant regions across the full time cycle (Pataky, 2010). In S1, we observed a marked increase in bilateral TA and GM muscle activation during stance in the early phase, which aligned with previous findings and likely reflected a rapid response to the altered walking conditions and the need to maintain dynamic stability (Dietz et al., 1994; MacLellan et al., 2014). SPM analysis further revealed shifts in both the timing and magnitude of bilateral ankle muscle activation, suggesting interlimb coordination for managing instability resulting from the unexpected perturbation introduced by the onset of split-belt walking. Specifically, the earlier onset of heightened bilateral TA and GM activity during weight acceptance in the early phase of S1 coincided with reductions in the first peak of the GRFs. This pattern of increased ankle co-contraction and decreased GRF at weight acceptance is suggestive of a cautious gait strategy at the onset of S1. By stiffening the joint with increased ankle muscle activity, participants may modulate weight acceptance by initially limiting body weight transfer onto the destabilizing fast leg (Ogawa et al., 2014). Elevated bilateral TA and GM activity were also observed during push-off in the early phase of S1, along with reductions in the second peak of GRFs. This is likely reflective of motor strategies favoring stability over generation of higher propulsive forces while sensorimotor prediction errors were still being recalibrated to split-belt walking. The use of SPM further reveals that both EMG and GRF patterns behaved similarly to baseline walking patterns by the late phase of S1 (specific to each belt’s baseline speed), which coincided with an improved SLS and a decline in EDA. Taken together, the observed motor adaptation patterns in EMG and GRFs further support the recalibration of SLS and demonstrate a co-modulation with physiological arousal during the first exposure to split-belt walking.

Repeated exposures to split-belt walking led to progressively attenuated responses across both the CNS and ANS. In S2 and S3, the magnitude of adaptation in SLS, muscle activity, GRFs and EDA was reduced compared to S1, demonstrating motor and physiological savings upon repeated exposure. Early-phase EDA levels in S2 and S3 dropped to approximately 40% of the individual’s session maximum. As gait parameters, such as SLS, stabilize and motor pattern refinements become more subtle, lower attentional demands are directed towards the task (Yoshida et al., 2024). Furthermore, this attenuation of physiological arousal may imply an improved perception of control or confidence in adapting to the task, as supported by lower RPS scores indicating one’s perception of improved walking balance stability in S2 and S3. Interestingly, physiological arousal consistently attenuates to around 20% of session maximum by the late phase of all three Split blocks, approaching levels seen during fast speed baseline walking. Similarly, mean SLS values in the late phases of S2 and S3 closely approximated each other, suggesting repeated exposure to split-belt walking may lead to a relatively stable gait pattern. Although speculative, the consistent late phase arousal levels seen in fast speed baseline walking and all split blocks may be due to a persistent ANS contribution necessary to meet ongoing physiological demands of the walking speeds, rather than a marker of task difficulty after behavioural adaptation plateaus (e.g., SLS reaches plateau). Similar patterns have been observed in postural and balance control tasks, where ANS-mediated arousal attenuated with repeated exposures to the task but remained somewhat elevated under persistent physical or cognitive load (Johnson et al., 2019; Zaback et al., 2019).

In-line with the attenuated physiological arousal observed in S2 and S3, patterns of muscle activation and GRFs also showed reduced magnitude of changes between early and late phases. Unlike S1, bilateral ankle muscle activity and kinetic adaptation patterns in S2 and S3 remained more consistent and exhibited less within-block variation. Although these patterns did not fully return to baseline walking levels, their relative stability in S2 and S3 may suggest that the motor system engaged in more efficient gait strategies with repeated practice. Energy efficiency is one of the primary goals of human walking, and our findings align with this principle (Kuo & Donelan, 2010; Selinger et al., 2015). While we did not directly measure metabolic cost, the reduction in muscle activation with improved SLS is consistent with prior work showing decreased energetic cost during a single bout of split-belt adaptation (Finley et al., 2013). Our results extend these insights by demonstrating that with repeated exposures, the motor system may not only recall prior adaptations but also appears to refine them, leading to more stable and efficient motor patterns.

Upon returning to tied-belt walking (i.e., de-adaptation), clear after-effects were observed in all measures. These after-effects indicate that the previously adapted motor patterns during split-belt walking temporarily persisted despite the reintroduction of symmetrical belt speeds (Malone et al., 2011; Reisman et al., 2005). However, these adapted patterns diminished quickly as the CNS reverted to the locomotor strategy suited for tied-belt walking, reflective of a well-established walking pattern. Among all measures, SLS showed significant changes between phases in T1 and T2 only, while all other measures demonstrated significant between-phase changes across all Tied blocks. Notably, EDA during the Early phase of each Tied block remained approximately 20% above baseline, which may reflect a carry-over from the arousal levels during the Late phase of the Split blocks (Fig 6). However, EDA levels during the Late phases of all Tied blocks returned to near levels of baseline slow walking. Taken together, phase-related changes in physiological arousal, bilateral muscle activity, and changes in GRFs all moved toward baseline slow walking levels, suggesting individuals re-familiarized with the symmetrical walking condition rapidly across both the CNS and ANS.

## Conclusions

Our study provides new insight into the co-modulation of the CNS and ANS responses during continuous walking challenges on a split-belt treadmill. Our findings offer a novel systems-level perspective on locomotor learning, supporting the co-modulation of the ANS alongside CNS motor adaptation to a continuous walking challenge. Future research should explore the underlying neural mechanisms driving this co-modulation and investigate how these processes differ in populations with motor impairments and/or decreased balance confidence. For example, individuals post-stroke have shown reduced attenuation of EDA and less motor adaptation in response to repeated postural perturbations compared to age-matched controls (Pollock et al., 2017). Understanding how CNS-ANS modulation in walking varies across populations may help inform more personalized approaches to gait rehabilitation.

## Authors’ Contributions

BL: Conceptualization, methodology, data analysis, writing – original draft, writing – review & editing

KJY: Conceptualization, methodology, data collection, data analysis, writing – review & editing

SJG: Methodology, writing – review & editing

TDI: Methodology, writing – review & editing

LAB: Conceptualization, writing – review & editing

CLP: Conceptualization, funding acquisition, methodology, supervision, writing – review & editing

## Availability of data and material

Data are available from the corresponding author upon reasonable request.

